# Dynamic functional adaptations during touch observation in autism: Connectivity strength is linked to attitudes towards social touch and social responsiveness

**DOI:** 10.1101/2024.09.30.615926

**Authors:** Haemy Lee Masson

**Author notes:** Correspondence should be addressed to Haemy Lee Masson Address: Department of Psychology, Durham University, South Road, Durham, DH1 3LE, UK.

## Abstract

Autistic adults often experience differences in social interactions involving physical contact. Brain imaging studies suggest that these differences may be related to atypical brain responses to social-affective cues, affecting both the experience of receiving touch and observing it in others. However, it remains unclear whether these atypical responses are limited to specific brain regions or represent broader alterations in brain connectivity. The current study investigated how the functional network architecture is modulated during touch observation associated with autism and explored the extent to which changes in this architecture are associated with individual differences in social touch preferences and social responsiveness. By integrating generalized psychophysiological interaction (gPPI) analysis with independent component analysis (ICA), the current study analyzed existing fMRI datasets, in which 21 autistic and 21 non-autistic male adults viewed videos of social and nonsocial touch while undergoing MRI scans. A gPPI analysis of pre-defined regions of interest revealed that autistic adults exhibited increased connectivity between sensory and social brain regions. The strength of some of these connections was positively associated with a higher preference for social touch and greater social responsiveness, suggesting neural compensatory mechanisms that may help autistic adults better understand the meaning of touch. At the level of large-scale brain networks extracted using ICA, atypical connectivity was predominantly observed between the sensorimotor network and other networks involved in social-emotional processing. Increased connectivity was observed in the sensorimotor network during nonsocial touch, suggesting that embodied simulation, the process by which individuals internally simulate touch experience of others in this context, may be more engaged when observing human-object interactions than during human-to-human touch in autism. This study reveals atypical context-dependent modulation of functional brain architecture associated with autism during touch observation, suggesting that challenges in recognizing and using affective touch in social interactions may be associated with altered brain connectivity. Neural compensatory mechanisms in autistic individuals who enjoy social touch and show higher social responsiveness may function as adaptive social responses. However, these compensations appear to be limited to specific brain regions, rather than occurring at the level of large-scale brain networks.

## Introduction

Touch plays a crucial role in social interaction (Field, 2010), effectively conveying social and emotional cues (Hertenstein et al., 2006a, 2006b, 2009; McIntyre et al., 2022) and fostering social bonds (Dunbar, 2010; Morrison et al., 2010; Suvilehto et al., 2015). Even when merely observing others’ social touch interactions, most of us accurately and rapidly grasp their meaning with high inter-observer reliability (Lee Masson and Op de Beeck, 2018). This process occurs in the brain within a few hundred milliseconds (Pihko et al., 2010; Schirmer and McGlone, 2018; Lee Masson and Isik, 2023) and involves a complex interplay of neural and cognitive processes (Ebisch et al., 2011a; Morrison et al., 2011; Bolognini et al., 2013; Lee Masson et al., 2018, 2020b; Schirmer and McGlone, 2018; Peled-Avron et al., 2019; Lee Masson and Isik, 2023). A recent study using Electroencephalography (EEG) and functional Magnetic Resonance Imaging (fMRI) revealed that the brain processes social-affective features of observed touch within 200 milliseconds through visual, social perceptual and somatosensory pathways (Lee Masson and Isik, 2023). Both meta-analysis study and literature review have provided solid evidence that observing social touch extensively involves the sensory cortex, the social-cognitive brain network, and the limbic system (Peled-Avron and Woolley, 2022; Schaefer et al., 2024).

Autism is a neurodevelopmental condition characterized by a wide range of cognitive, sensory, and behavioral differences (American Psychiatric Association, 2013). While social touch in autism remains relatively underexplored, touch is a significant aspect of daily life for autistic individuals and is often influenced by their sensory differences (O. Miguel et al., 2017; Ujiie and Takahashi, 2022). This can include distinctive patterns of touch sensitivity, such as heightened or reduced responsiveness to tactile stimuli, as well as particular behaviors related to touch, including social touch avoidance (Baranek et al., 1997; Baranek, 1999; Cascio et al., 2012, 2016; Ludlow et al., 2015; Mammen et al., 2015; Mikkelsen et al., 2018; Lee Masson et al., 2019, 2020a; Mello et al., 2024). While much of the previous research has centered on tactile reactivity and its possible influence on social touch avoidance, a recent study explored cases where autistic children resist social touch from their parents and highlighted that the touch avoidance might be attributed not to tactile reactivity, but to the social nature of the touch interfering with the child’s ongoing activities (Henderson, 2022). Research on touch observation has also documented atypical processing of visually presented touch associated with autism. Adults with high autistic traits rate visually presented gentle stroking touch as less pleasant than those with low autistic traits (Haggarty et al., 2021). Autistic adults (AUT) exert more cognitive effort, as indicated by larger pupil dilation, when observing social touch compared to non-autistic adults (NON-AUT) (Lee et al., 2024). These behavioral and physiological differences are also reflected in the brain. AUT exhibit atypical neural responses to images of social touch interactions, as evidenced by increased activity in both early sensory (P1) and later high-order cognitive (Late Positive Potential) neural components (Peled-Avron and Shamay-Tsoory, 2017) and atypical somatosensory neural response patterns (Lee Masson et al., 2019). The above evidence suggests that differences may arise not in a single cognitive or neural domain but in multiple systems and their interactions.

Atypical neural connectivity has been suggested as a core neural marker of autism (Just et al., 2004, 2007; Lewis et al., 2014, 2017). Numerous studies, including those cited above, have reported widespread underconnectivity, particularly in long-range connections. This includes interhemispheric connectivity, connections between the anterior and posterior components of the default mode network (DMN), the insula, and the frontal areas with other cortical regions (Martino et al., 2014; Cheng et al., 2015; Abbott et al., 2016; O’Reilly et al., 2017). Remarkably, reduced network efficiencies in the white matter structures associated with low-level sensory processing has even been observed in six-month-old infants who were later diagnosed with autism (Lewis et al., 2017). Reduced fractional anisotropy and increased radial diffusivity have also been reported, suggesting smaller axon diameters and poor myelination, potentially leanding to inefficient communication between brain regions (Zikopoulos and Barbas, 2013). However, more recent studies testing the underconnectivity hypothesis of autism present mixed results (Maximo et al., 2014; Hahamy et al., 2015; Picci et al., 2016; Liloia et al., 2024), with evidence of overconnectivity between the DMN and the executive control network (ECN) (Abbott et al., 2016). However, most of the studies above have adopted a task-free resting state or structural connectivity approach to gain insight into the intrinsic (baseline) brain connectivity of autism.

Task-based functional connectivity (FC) is crucial for understanding how extensive brain networks coordinate to support complex cognitive functions (Cohen and D’Esposito, 2016; Gonzalez-Castillo and Bandettini, 2018). This approach is particularly relevant for social touch research, as observing social touch leads to the increased functional communication of multiple neural systems, from early sensory processing to high-level social cognitive systems (Lee Masson et al., 2020b). However, in autism, it remains unclear how observing social versus nonsocial touch affects the communication between these networks and how attitudes towards social touch and autistic social traits are linked to functional connectivity.

The present study investigates how brain regions and networks involved in various perceptual and cognitive functions coordinate to support task demands during the observation of social and nonsocial touch in autistic adults. Additionally, it examines whether attitudes towards social touch and autistic social traits are linked to the functional coordination of brain networks. This study reanalyzed the dataset from the previous work (Lee Masson et al., 2019) to investigate the functional modulation of brain connectivity during touch observation. This research question was not addressed in the original publication. The study aim was operationalized by applying generalized psychophysiological interactions (gPPI) and independent component analysis (ICA) to functional magnetic resonance imaging (fMRI) data from participants who viewed social and nonsocial videos. gPPI examines how connectivity among brain regions and networks changes in response to different psychological contexts (McLaren et al., 2012). ICA decomposes complex fMRI data into independent spatial and temporal components, allowing for the identification of distinct brain networks during touch observation (Mckeown et al., 1998). This combined approach illuminates how different types of touch (social versus nonsocial) modulates brain connectivity in autism, an area that has not been previously explored. Based on prior findings of differences in sensory cortical responses and high-level cognitive neural responses during social touch observation, as discussed above, functional communication between brain areas and networks involved in various perceptual and cognitive functions may differ in autism. Given the mixed findings regarding under- and over-connectivity discussed earlier, this study refrained from hypothesizing whether the connectivity would be reduced or enhanced.

## Methods

To address current research questions using a gPPI analysis approach, existing fMRI data was re-analyzed (Lee Masson et al., 2019). While the original study investigated how neural patterns of individual brain regions reflect the social-affective dimension of observed touch through representational similarity analysis (RSA), the current study examines the changes in connectivity strength during the observation of social versus nonsocial touch. The FC methods offer unique insights that may not correspond with the findings from RSA (Pillet et al., 2020).

### Participants

The sample size of the current study is identical to the original study (Lee Masson et al., 2019). The original study includes functional and anatomical MRI scans and behavioral measures from 42 male participants, 21 of whom were NON-AUT and 21 of whom had previously received an autism diagnosis via a multidisciplinary team at the Expertise Center for Autism at the University Hospitals Leuven, following DSM-IV or DSM-5 criteria. Both groups were matched on sex (all male), age (mean age: 25 for AUT and 23.9 for NON-AUT), and IQ (111.3 for AUT and 111.5 for NON-AUT). All inclusion and exclusion criteria were established in the original study. All participants provided written informed consent, and the Medical Ethical Committee of KU Leuven approved the original study (S53768 and S59577).

### Experiment

Forty-two male participants watched 39 three-second-long social (e.g., hugging a person) and 36 nonsocial video clips (e.g., carrying a box) during fMRI scans in the original study. Figure 1 shows examples of social and nonsocial touch. The complete set of video stimuli can be found in the prior work (Lee Masson and Op de Beeck, 2018). Notably, visual features such as luminance, motion energy, and biological motion are matched between social and nonsocial touch videos.

**Figure 1.**
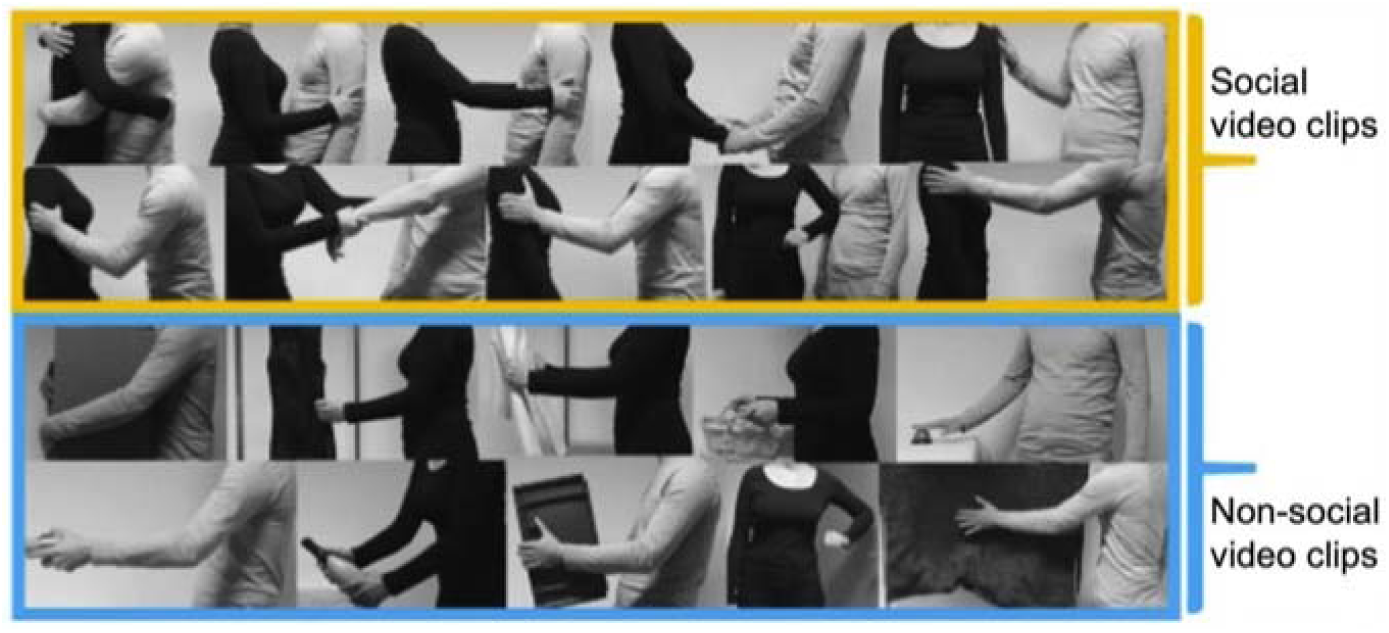
Example frames from video clips. The images in the yellow box show frames from social touch videos, which feature both pleasant and unpleasant touch interactions such as hugging, caressing, handholding, and slapping. Meanwhile, the images in the blue box are from nonsocial touch videos, displaying matched human-object interactions. This figure is published under a CC-BY-NC-ND license (https://creativecommons.org/licenses/by-nc-nd/4.0/) and is reused from Figure 1 in the original study (Lee Masson et al., 2018). The complete set of original video materials is available at https://osf.io/8j74m/.

All participants completed two self-report assessments of social touch and social responsiveness. The Social Touch Questionnaire (STQ) measures an individual’s attitude toward social touch using a 5-point Likert scale (1 – strongly disagree, 5 – strongly agree), with lower scores reflecting a greater tendency to avoid social touch (Wilhelm et al., 2001). The STQ consists of 20 items, with an example item being, “I generally like when people express their affection towards me in a physical way.” The social responsiveness scale (SRS) is a standardized instrument designed to measure social differences in individuals using a 4-point Likert scale (1 – not true, 4 – always almost true), with higher scores reflecting greater social differences (Constantino et al., 2003). The SRS comprises 65 items that assess social awareness, social cognition, social communication, social motivation, and autistic mannerisms. An example item is: “I feel much more uncomfortable in social situations than when I am alone.”

### MRI acquisition, preprocessing, and head motion

Whole-brain images (37 slices, a voxel size of 2.7 × 2.7 × 3 mm3) were obtained on a 3T Philips scanner with a 32-channel coil and an echo-planar (EPI) T2∗-weighted sequence. The acquisition parameters were as follows: repetition time (TR) = 2000 ms, echo time (TE) = 30 ms, flip angle (FA) = 90°, and field of view (FOV) = 216 × 216 mm^2^. The source study involved an average of seven functional runs with each run comprising of 239 volumes, resulting in a total acquisition of 1673 volumes for the main experiment per participant. Structural MR images were collected using a T1-weighted sagittal high-resolution magnetization-prepared rapid gradient echo (MPRAGE) sequence. The acquisition parameters were as follows: TR□=□9.6 ms, TE□=□4.6□ms, FA□=□8°, FOV□=□250□×□250 mm^2^, voxel size□=□0.98□×□0.98□×□1.2□mm, 182 axial slices.

The current study used preprocessed fMRI data from the source study. The preprocessing steps included slice timing correction, realignment of functional images to the mean image of the first run, normalization to a Montreal Neurological Institute (MNI) space with a re-sampling size of 3 × 3 × 3 mm, and spatial smoothing with an 8 mm full-width at half maximum Gaussian kernel. Smoothed preprocessing data was used for ICA, while unsmoothed preprocessing data was used for connectivity analysis to avoid spillage effects (Alakörkkö et al., 2017). Importantly, both groups demonstrated similar levels of head motion. There were no significant differences in maximum head motion (AUT: average 1.38 mm, NON-AUT: 1.36 mm, *t*(40) = 0.04, *p* = 0.97) or in mean frame-wise head motion displacement (AUT: average 0.13 mm, NON-AUT: 0.13 mm, *t*(40) = 0.04, *p* = 0.96).

### Identifying brain regions processing observed touch

Most regions of interest (ROIs) are predefined in the source study based on their roles in touch observation. Brodmann Area (BA) 17, 18, 19, 37, and V5 were selected as they process visual information (Sunaert et al., 1999; Wohlschläger et al., 2005; Thompson and Baccus, 2012). Middle temporal gyrus (MTG), superior temporal gyrus (STG), precuneus (Precu), and temporoparietal Junction (TPJ) were included as they are implicated in social processing (Jacoby et al., 2016). These ROIs were functionally selected from the main fMRI runs with the contrast of touch observation > fixation cross. Lastly, BA1, 2, 3 in the somatosensory cortex were included as they process tactile information (Kaas, 1983). These areas were extracted from the localizer run where participants received actual affective touch (pleasant and unpleasant > rest). All functional ROIs were defined based on activation within each anatomical template, using the second-level group results (N=42, encompassing both groups). ROIs are shown in Figure 2. The same set of ROIs was applied to both groups, as the source study found no group differences in brain activation. For more details, refer to the source study (Lee Masson et al., 2019).

**Figure 2.**
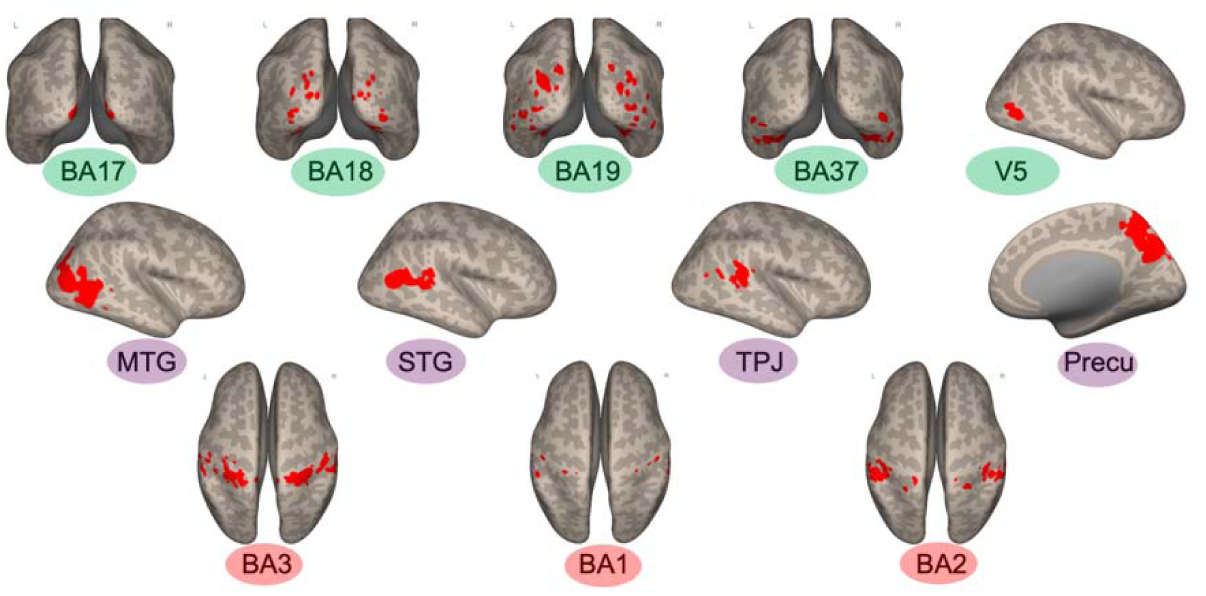
Visualization of the selected ROIs. The red markings on the brain images indicate the regions corresponding to each ROI. Visual areas are denoted by the color green, social brain regions are indicated by the color purple, and somatosensory areas are represented by the color red.

### Identifying brain networks processing observed touch

ICA is a data-driven multivariate approach that does not rely on prior assumptions about the brain regions involved in touch observation. ICA method combined with connectivity analysis was used to comprehensively measure brain network communication, complementing ROI-based analysis. Specifically, spatial ICA, implemented in the Group ICA Toolbox (GIFT version 4.0.4.11), decomposes whole-brain fMRI data and extracts spatially independent components (ICs) (Rachakonda et al., 2007). For this analysis, the preprocessed fMRI data from the source study was entered into GIFT to identify groups of brain regions that have temporally coherent BOLD signal fluctuations during touch observation.

Similar to prior studies that employed ICA (Cisler et al., 2013; Jung et al., 2019; Lee Masson et al., 2020b, 2020a), the optimal number of ICs that could accurately account for the total variance of the fMRI data was determined using the minimum description length (MDL) criterion (Li et al., 2007). The optimal number of ICs was determined to be 28. The dimensionality of the fMRI data was reduced using standard principal components analysis (PCA) at the individual and group level before ICA was performed. To ensure stability, the Infomax algorithm was used and repeated ten times with the ICASSO toolbox implemented in GIFT. This repetition enabled the extraction of the 28 most reliable and stable ICs at the group level (Bell and Sejnowski, 1995). For the subject-level ICs, GICA back-reconstruction was performed on the group ICs (Calhoun et al., 2001). The spatial images and time-courses were transformed into standardized z-scores. The z-score of each voxel represents its contribution to the time course of each IC. As a final step, subject-level ICs were used to create a group mean spatial map and a group mean time course for each IC.

Task-relevant ICs were selected from 28 with spatial and temporal sorting implemented in GIFT, using methods identical to the prior work (Lee Masson et al., 2020b). Nine out of the 28 ICs were related to artifacts: six were located outside of the grey matter (GM), two were in the cerebrospinal fluid (CSF), and one was identified as the cerebellum, which was not fully scanned in the source study. The remaining 19 ICs were labeled based on the spatial correlation between each IC and the brain template included in GIFT (Smith et al., 2009). The label was selected based on the template that has the highest correlation with the IC (Table1). These ICs will be referred to as network based on the above results (e.g., higher visual network instead of IC1). Some network labels were named according to the brain regions they include. Lastly, the temporal sorting method was used to measure the degree of synchronization (task-relatedness) between the time course of the network and stimulus events for social, non-social, and baseline conditions (Calhoun et al., 2008). Networks that showed statistically different degrees of synchronization based on stimuli conditions were selected for further investigation.

The final set of brain networks included in the main connectivity analysis consisted of the higher visual network, the left executive control network, the sensorimotor network, the social perceptual network, the posterior and anterior salience networks, the limbic system, the reward system, and two default mode networks (networks marked with an asterisk (*) in Table 1). Using SPM12 (Wellcome Department of Imaging Neuroscience, UCL, London, United Kingdom) in MATLAB R2020a (The MathWorks Inc., Natick, MA, United Kingdom), the binary map of each selected network was generated by conducting a one-sample t-test with a threshold of family-wise error (FWE) corrected P-value less than 0.001 (Figure 3). Table S1 in the Supplementary Materials provides the peak x, y, z coordinates, the names of the brain areas, and the number of voxels included in each network. Notably, the two-sample t-test yielded no significant group difference in the degree of contribution of each voxel on the network (*p _FWE_* < 0.05), indicating that the NON-AUT and AUT group did not differ in how the functional brain networks formed during touch observation.

**Figure 3.**
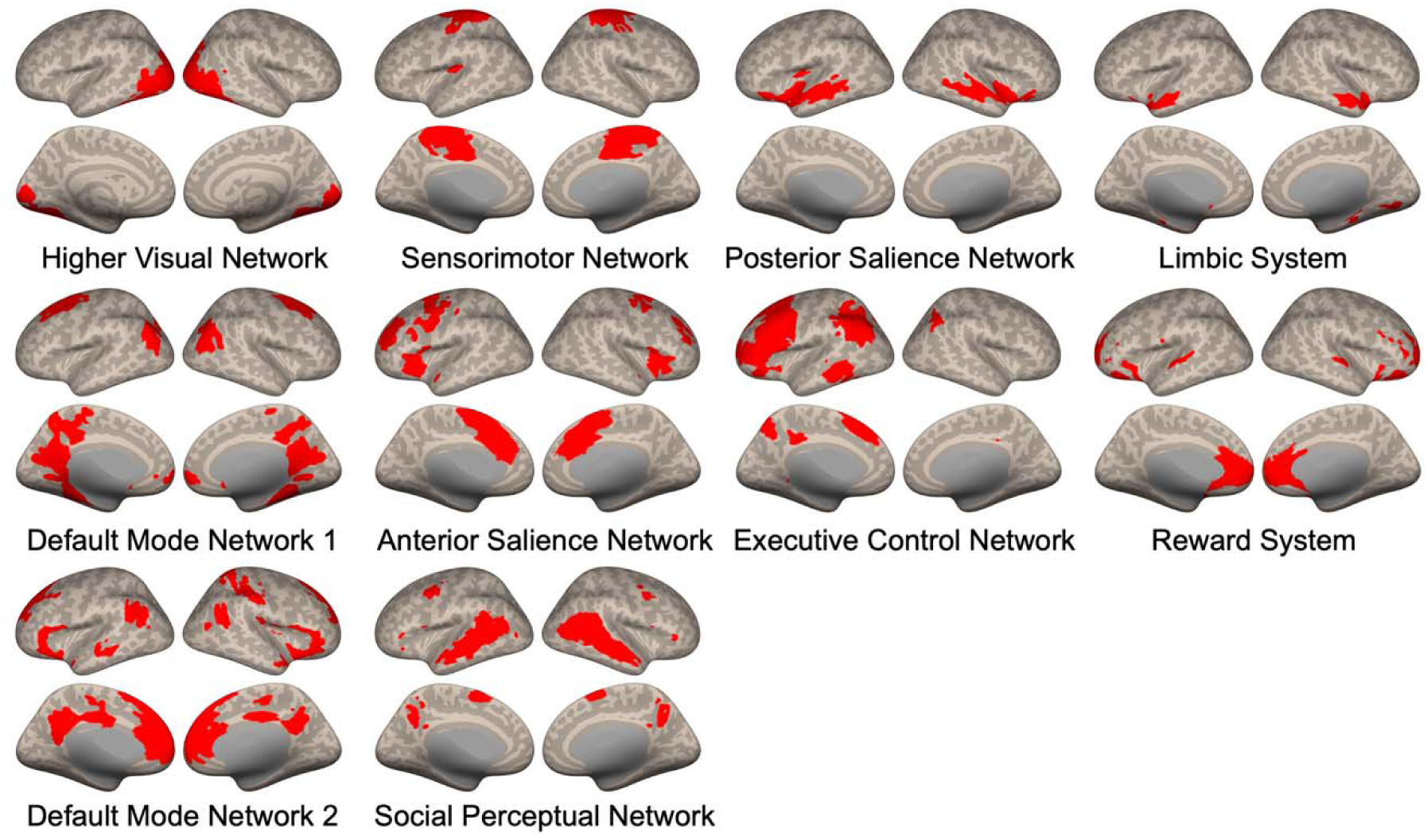
Ten task-relevant brain networks, identified by ICA. The red markings on the brain images indicate the regions included in each network.

**Table 1.** Labels for each component. Labels are determined by the spatial correlation (*r*) between the selected template providing the label and the group averaged spatial map of each component. When a component is situated outside of gray matter, it is categorized as noise. When a component is task-relevant according to the temporal sorting methods, it is indicated with an asterisk (*).

| Component Number | Component Description | $r$ |
| --- | --- | --- |
| IC1 | Higher Visual Network (*) | 0.21 |
| IC2 | Noise |  |
| IC3 | Precuneus Network | 0.37 |
| IC4 | Sensorimotor Network (*) | 0.16 |
| IC5 | Primary Visual Network | 0.27 |
| IC6 | Noise |  |
| IC7 | Noise |  |
| IC8 | Noise |  |
| IC9 | Ventral Default Mode Network | 0.45 |
| IC10 | Noise |  |
| IC11 | Posterior Salience Network (*) | 0.16 |
| IC12 | Auditory Network | 0.29 |
| IC13 | Noise |  |
| IC14 | Sensorimotor Network | 0.33 |
| IC15 | Noise |  |
| IC16 | Right Executive Control Network | 0.30 |
| IC17 | Ventral Default Mode Network | 0.18 |
| IC18 | Dorsal Default Mode Network (*)/Limbic System | 0.10 |
| IC19 | Dorsal Default Mode Network/DMN1 (*) | 0.33 |
| IC20 | CSF |  |
| IC21 | Cerebellum |  |
| IC22 | CSF |  |
| IC23 | Visuospatial Network | 0.14 |
| IC24 | Anterior Salience Network (*) | 0.27 |
| IC25 | Left Executive Control Network (*) | 0.20 |
| IC26 | Dorsal Default Mode Network (*)/Reward System | 0.14 |
| IC27 | Dorsal Default Mode Network/ DMN2 (*) | 0.49 |
| IC28 | Language Network (*)/Social Perceptual Network | 0.20 |

### FC between ROIs and networks

Additional preprocessing and FC analysis were performed with CONN toolbox (version 22.a) (Whitfield-Gabrieli and Nieto-Castanon, 2012; Nieto-Castanon and Whitfield-Gabrieli, 2021). Prior to gPPI analysis, the artifact detection and repair toolbox implemented in CONN was used to remove outlying volumes with framewise displacement above 0.9 mm or global BOLD signal changes above 5 standard deviations. A standard denoising pipeline was applied to the functional data. This involved regressing out white matter (WM) and CSF timeseries, motion parameters, artifact-related covariates, outlying volumes, session and task effects, and linear trends within each functional run. A high-pass filter was applied to eliminate slowly fluctuating signals, such as scanner drift, at 0.008 Hz.

gPPI method was used to measure the changes in FC across social and nonsocial touch conditions. ROI-ROI connectivity analysis includes 12 ROIs implicated in touch observation – 5 visual, 4 social, and 3 somatosensory areas. Network-network connectivity analysis includes 10 large scale brain networks identified with ICA. The analysis of ROIs and network FC was conducted separately. In the subject-level analysis conducted in the CONN toolbox, a gPPI model was created for each pair of seed and target ROIs/networks. This model included seed blood oxygenation level dependent (BOLD) signals as physiological factors, boxcar signals representing each touch condition (convolved with an SPM canonical hemodynamic response function) as psychological factors, and their interaction as psychophysiological interaction terms. Changes in connectivity strength across touch conditions were evaluated by the regression coefficient of the psychophysiological interaction terms. This coefficient measured how well the interaction terms explained variations in the target BOLD signals as the dependent variable (McLaren et al., 2012).

Group-level analyses were performed using generalized linear mixed-effects models (GLM) implemented in R (R Core Team, 2024) with the lme4 package (Bates et al., 2015). In the GLM analysis, connectivity measures for each touch condition from the subject-level analysis became the dependent variable. The model included touch condition (social vs. nonsocial touch), group (AUT vs. NON-AUT), STQ and SRS scores, and all possible interactions among these variables as independent variables. Random effects were also accounted for across subjects. Subsequently, with the car package (Fox and Weisberg, 2019), a mixed-model repeated-measures analysis of variance (ANOVA) was conducted to evaluate the significance of the effects of the independent variables on the GLM fit outcome. To address multiple comparisons involving tests on 66 ROI pairs and 45 network pairs, the p-values were adjusted using false discovery rate (FDR) correction. When the ANOVA results indicated a significant effect of the group or an interaction involving the group and other independent variables, a post hoc Tukey’s honest significant difference test was conducted using the emmeans package (Lenth, 2024) and reported in the results section. Since this study employed mixed models, the Kenward-Roger method was applied to precisely estimate the degrees of freedom by accounting for both the fixed and random effects in the model. Effect size (Cohen’s d) and confidence interval (CI) were calculated with the emmeans package using eff_size function. Spearman’s rank correlation followed by the FDR correction was used to evaluate the relationship between FC strength and both STQ and SRS scores, when interaction effects were found. The main effect of touch condition – viewing videos showing social versus nonsocial touch events – is not reported in the current study, as its impact on FC changes was documented in a prior study with a larger sample size of NON-AUT participants (Lee Masson et al., 2020b).

## Results

### The effect of group, touch type, social touch avoidance, and social responsiveness on the connectivity strength between brain regions

Out of the 66 ROI pairs, four showed a significant group effect or an interaction effect involving the group and other independent variables on FC strength. These effects were observed in the FC between BA18 and BA2, BA19 and Precu, V5 and BA1, and BA17 and STG. Specifically, ANOVA results revealed that the group factor significantly affects the FC strength between the early visual area (BA18) and the early somatosensory area (BA2) (*_χ_2*(1, 42) = 7.53, *p _FDR_* = 0.045). Post hoc tests revealed that the AUT group showed significantly higher FC strength compared to the NON-AUT group across two touch types (overall: *t*(34) = 2.76, *p* = 0.009, Cohen’s *d* = 3.37, CI = [0.82 – 5.92]; nonsocial: *t*(37.79) = 2.72, *p* = 0.0097, Cohen’s *d* = 3.42, CI = [0.81 – 6.03]; social: *t*(37.79) = 2.65, *p* = 0.01, Cohen’s *d* = 3.33, CI = [0.72 – 5.93]). Both groups show stronger FC during social touch compared to nonsocial touch (AUT: *t*(34) = −2.49, *p* = 0.02, Cohen’s *d* = −0.90, CI = [−1.64 – −0.15]; NON-AUT: *t*(34) = −2.18, *p* = 0.04, Cohen’s *d* = −0.99, CI = [−1.92 – −0.05]). Figure 4A illustrates the FC strength across groups and touch types.

**Figure 4.**
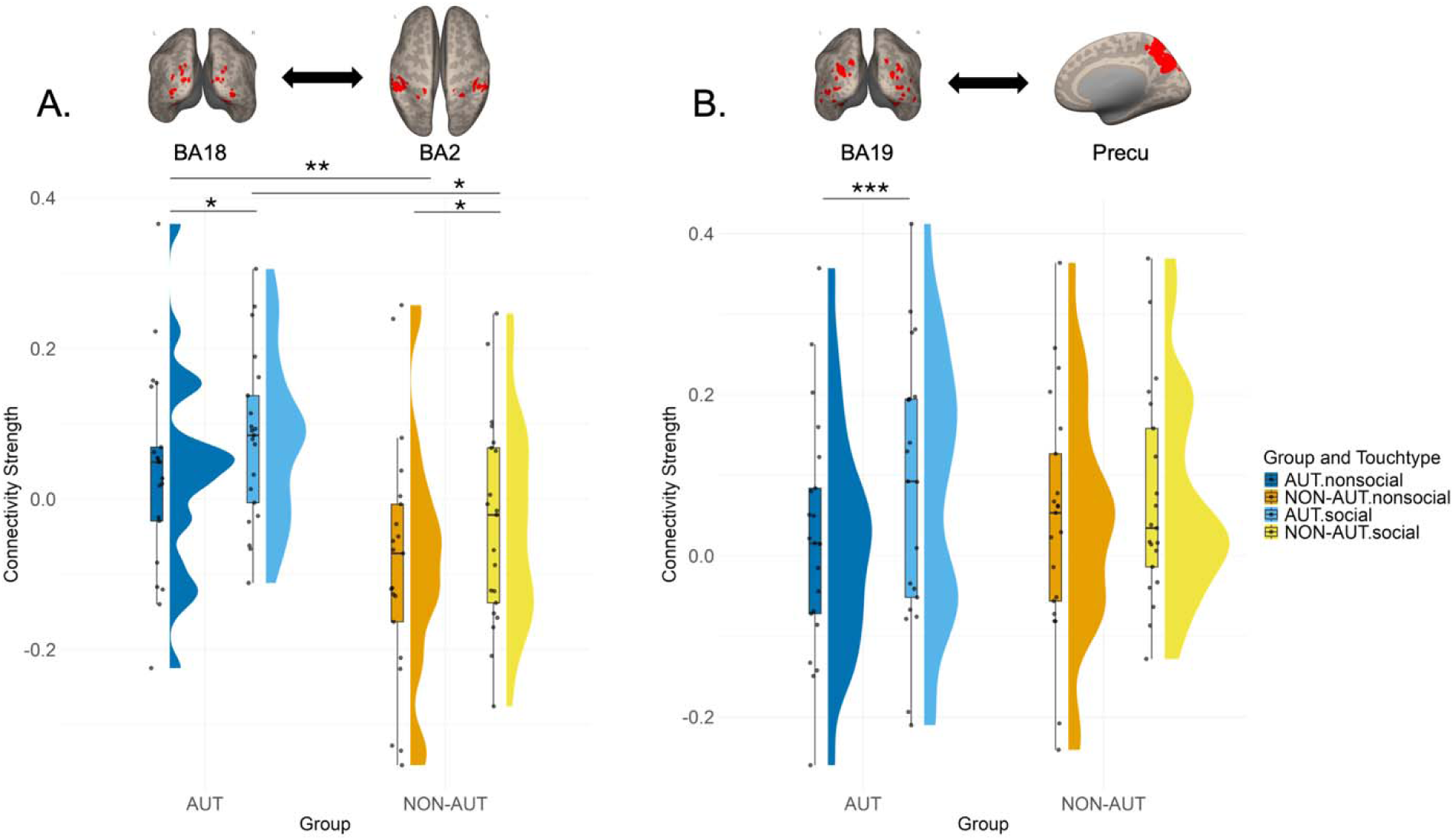
Connectivity strength across groups and touch types. **A.** FC strength between BA18 and BA2. **B.** FC strength between BA19 and Precu. The Y-axis shows FC strength between a selected ROI/network pair. The boxplots illustrate the median (a central line), interquartile range (the box representing the first to third quartiles), overall data spread, and outliers (points outside the whiskers). Individual data points are plotted on the boxplots. The half-eye plots adjacent to each boxplot illustrate the distribution shape of connectivity strength, highlighting where data points are concentrated (peaks). AUT data are colored in shades of blue (nonsocial in blue, social in sky blue), while NON-AUT data are colored in shades of orange (nonsocial in orange, social in yellow). Asterisks (*) are used to indicate levels of statistical significance: *: *p* < 0.05, **: *p* < 0.01, ***: *p* < 0.001.

Likewise, a significant interaction effect was found between group, touch type, and SRS scores on the FC strength between the visual area (BA19) and the social brain region (Precu) (*_χ_2*(1, 42) = 8.97, *p _FDR_* = 0.02). The AUT group showed a significant difference in FC strength between social and nonsocial touch, with social touch resulting in higher FC strength (*t*(34) = −4.09, *p* = 0.0002, Cohen’s *d* = −1.47, CI = [−2.25 – −0.70]). The NON-AUT group did not show a significant difference between the two touch types (Figure 4B). Interestingly, Spearman correlation analysis revealed that in the AUT group, there was a significant negative relationship between FC strength and SRS scores for both social (*_ρ_* = −0.68, *p _FDR_* = 0.0006) and nonsocial touch (*_ρ_* = −0.49, *p _FDR_* = 0.02), with a stronger effect observed in social touch (Figure 5A). AUT with greater social responsiveness (lower SRS scores) exhibited stronger FC between BA19 and Precu during social touch observation. In contrast, the correlations in the NON-AUT group were not significant, suggesting that social responsiveness was not linked to the connectivity strength between BA19 and Precu in these individuals.

**Figure 5.**
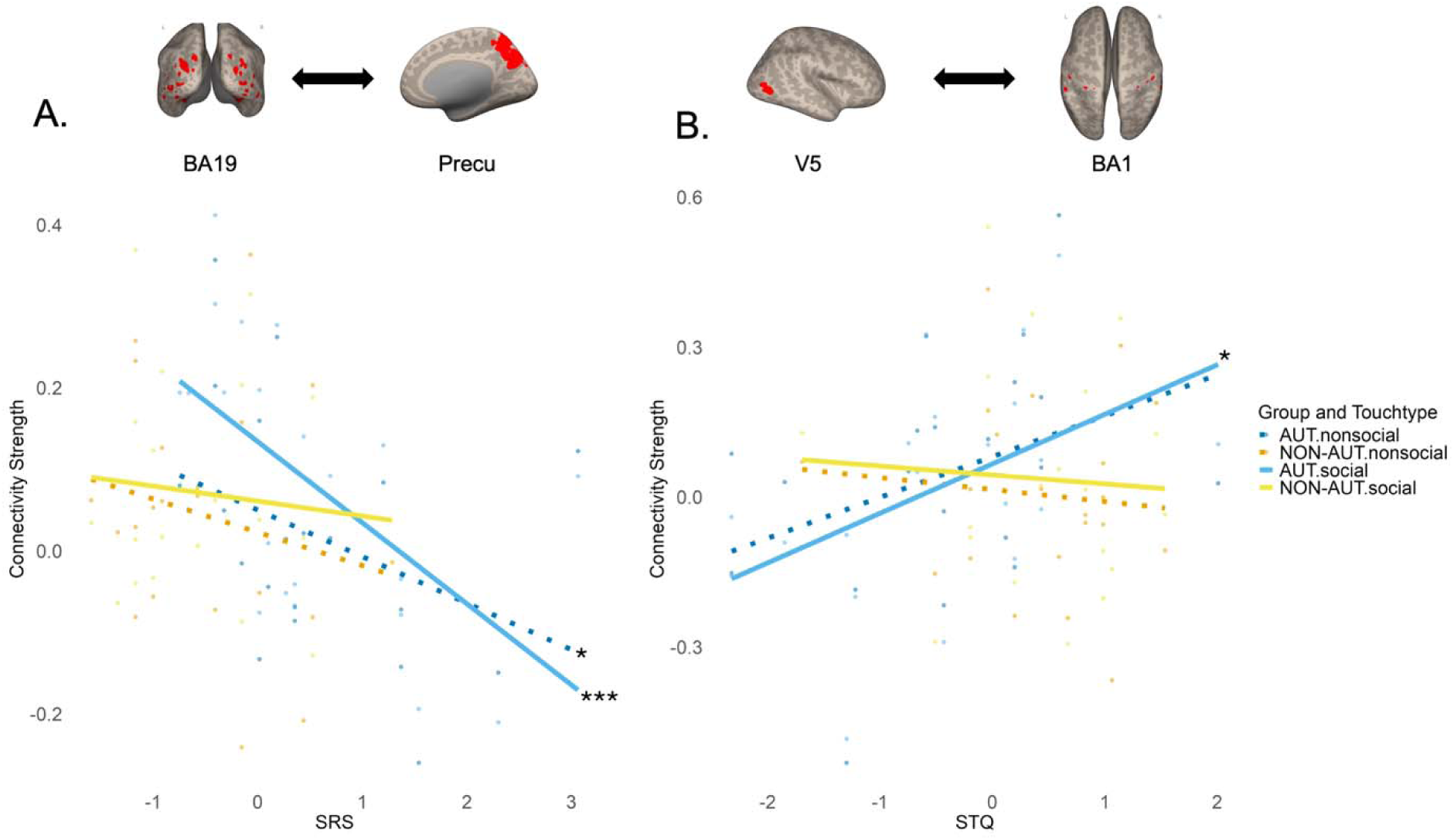
The relationship between connectivity strength and questionnaire scores. **A.** Relationship between FC strength of BA19 - Precu pair and SRS scores. **B.** Relationship between FC strength of V5 - BA1 pair and STQ scores. A higher SRS score indicates poorer social responsiveness, while a higher STQ score reflects a more positive attitude towards social touch. The plots illustrate the relationship between questionnaire scores (x-axis) and connectivity strength (y-axis) across different groups and touch types. Individual data points are displayed, with linear regression lines illustrating the direction and strength of the relationship. AUT data are shown in shades of blue (nonsocial in blue, social in sky blue), while NON-AUT data are shown in shades of orange (nonsocial in orange, social in yellow). Social touch is shown with a solid line, while nonsocial touch is shown with a dotted line. Asterisks (*) next to the right end of the regression lines denote the levels of statistical significance: *: *p* < 0.05, **: *p* < 0.01, ***: *p* < 0.001

A significant interaction effect was found between group and STQ scores on the FC strength between the visual motion area (V5) and the early somatosensory region (BA1) (*_χ_2*(1, 42) = 9.47, *p _FDR_* = 0.03). FC strength did not differ across groups and touch types. Yet, Spearman correlation analysis revealed that, in the AUT group, there was a positive correlation between STQ scores and FC strength during social touch observation (*_ρ_* = 0.53, *p _FDR_* = 0.01) and a marginal trend of positive correlation during nonsocial touch (*_ρ_* = 0.41, *p _FDR_* = 0.07). AUT with a positive attitude towards social touch (higher STQ scores) exhibited stronger FC between V5 and BA1. In contrast, in the NON-AUT group, the correlation was not significant, suggesting that the attitude towards social touch was not linked to FC strength in these individuals (Figure 5B).

Lastly, A significant interaction effect was found between group, STQ, and SRS scores on the FC strength between the early visual area (BA17) and the social brain region (STG) (*_χ_2*(1, 42) = 9.05, *p _FDR_* = 0.04). However, FC strength did not differ across groups and touch types. Likewise, correlation analysis showed no relationship between FC strength and the questionnaire scores.

### The effect of group, touch type, social touch avoidance, and social responsiveness on the connectivity strength between the large-scale brain networks

Out of the 45 network pairs, four demonstrated a significant group effect or an interaction effect involving the group and other independent variables on FC strength. These effects were observed in the FC between the sensorimotor network and posterior salience network, the sensorimotor network and DMN2, the executive control network and social perceptual network, and the limbic system and social perceptual network. Specifically, the FC strength between the sensorimotor network and posterior salience network varied significantly by touch type between the AUT and NON-AUT groups, as revealed by the significant interaction effect (*_χ_2*(1, 42) = 9.73, *p _FDR_* = 0.03). In the AUT group, FC strength between these networks was significantly higher during nonsocial touch compared to social touch (*t*(34) = 2.16, *p* = 0.04, Cohen’s *d* = 0.78, CI = [0.034 – 1.52]). In contrast, the NON-AUT group showed significantly lower FC strength during nonsocial touch compared to social touch (*t*(34) = −2.26, *p* = 0.03, Cohen’s *d* = −1.02, CI = [−1.96 – −0.09]). The effect of touch type on FC strength was modulated by group, with AUT individuals showing stronger connectivity in nonsocial touch and NON-AUT individuals showing stronger connectivity in social touch (Figure 6).

**Figure 6.**
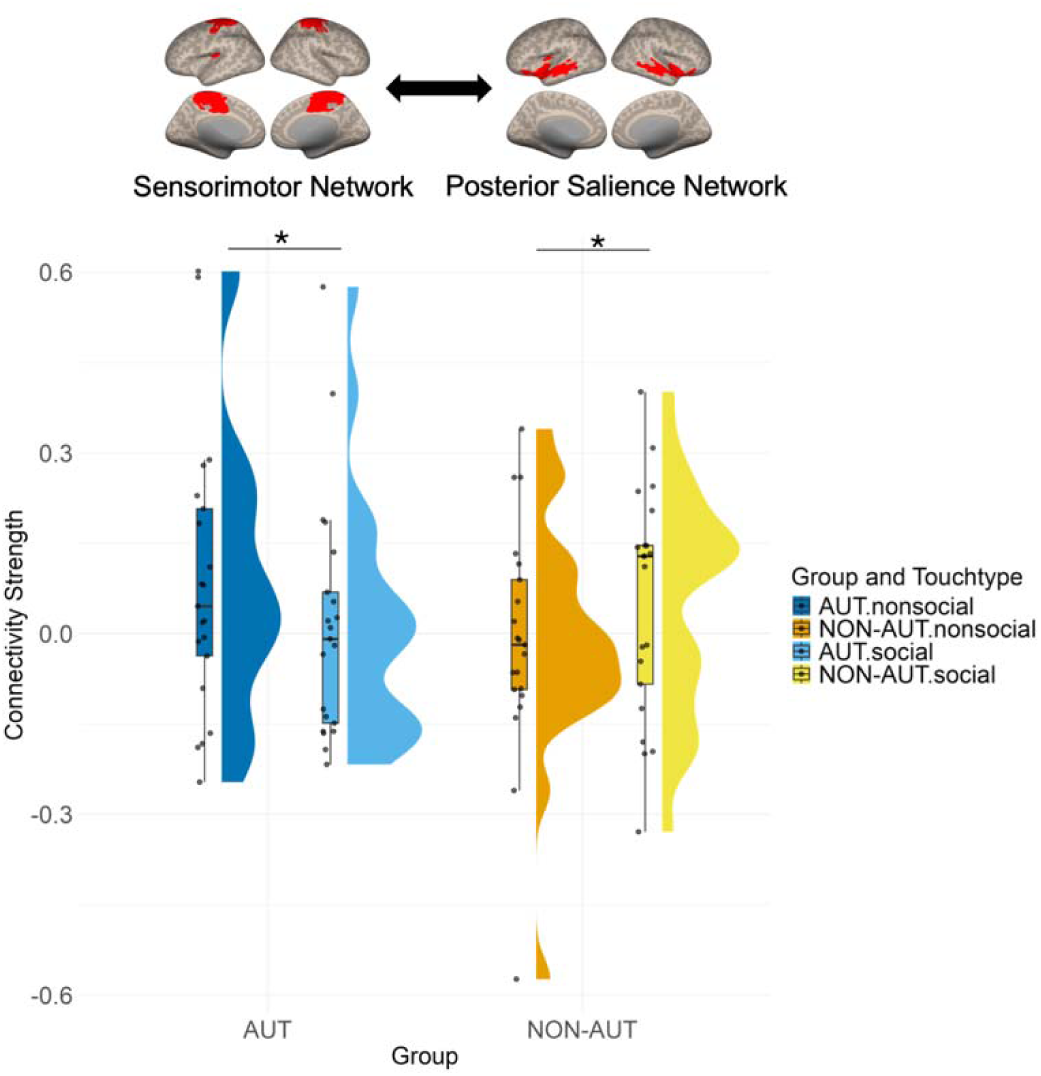
Connectivity strength between sensorimotor network and posterior salience network across groups and touch types. The plotting conventions are identical to those used in Figure 4.

Two groups showed differences in how STQ scores affected FC between the sensorimotor network and DMN2. The ANOVA revealed a significant interaction effect between group, touch type, and STQ scores on FC strength (*_χ_2*(1, 42) = 18.27, *p _FDR_* = 0.0002). This effect was mainly driven by a significant positive correlation between STQ scores and FC strength (*_ρ_* = 0.50, *p _FDR_*= 0.02) in the NON-AUT group during social touch observation (Figure 7A). Meanwhile, in the AUT group, the correlations are not significant during either nonsocial or social touch observation, suggesting no relationship between social touch preferences and FC strength. FC strength between these networks did not differ across groups and touch types.

**Figure 7.**
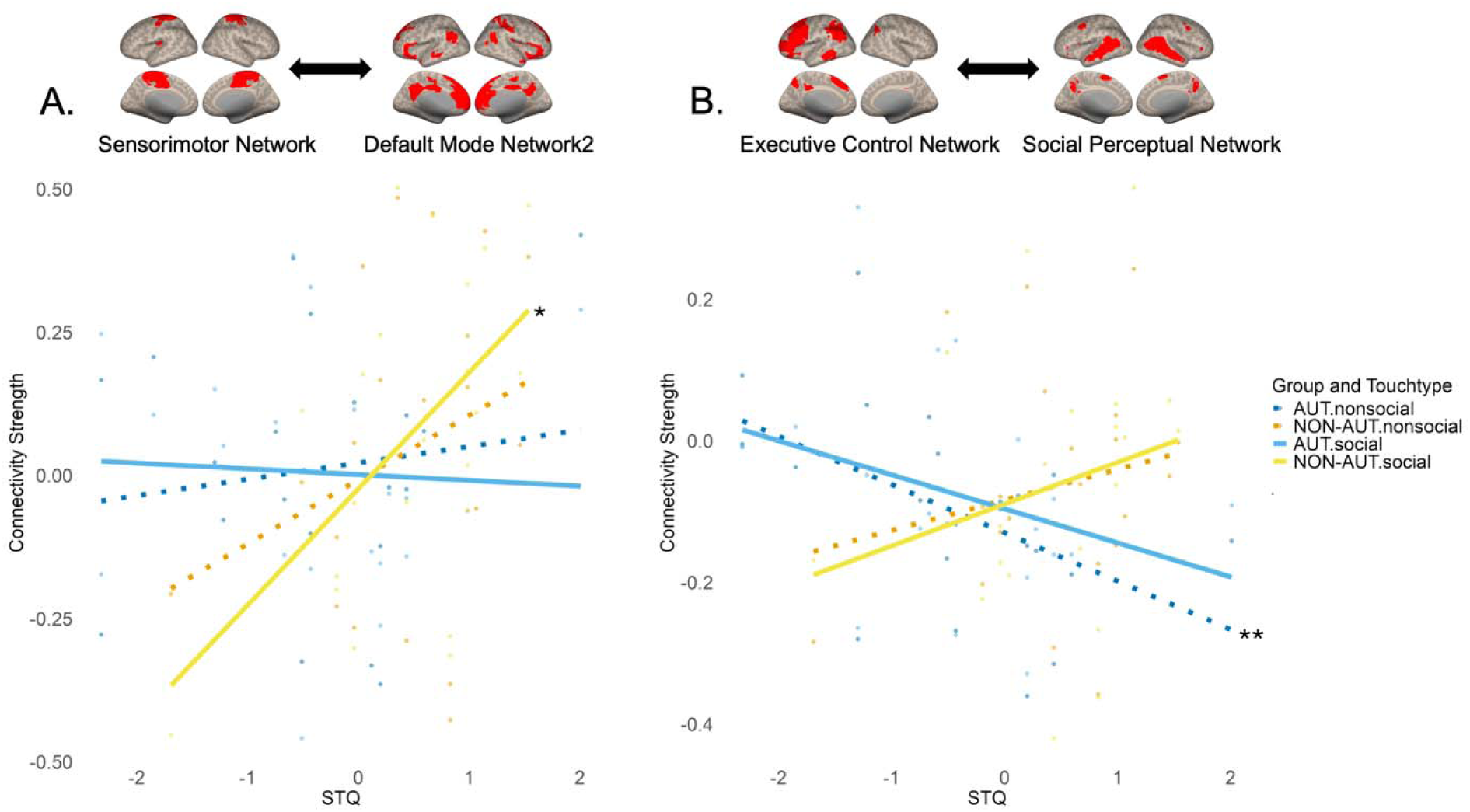
The relationship between connectivity strength and STQ scores. **A.** Relationship between FC strength of sensorimotor network – DMN2 pair and STQ scores. **B.** Relationship between FC strength of executive control network – social perceptual network pair and STQ scores. The plotting conventions are identical to those used in Figure 5.

The combined effect of STQ and SRS scores on FC strength between the executive control network and the social perceptual network varied across groups and touch types. ANOVA results showed a significant interaction between group, touch type, and STQ and SRS scores affecting FC strength (*_χ_2*(1, 42) = 9.08, *p _FDR_* = 0.04). This effect was primarily driven by the AUT group, which showed a significant negative correlation between STQ scores and FC strength in nonsocial touch (*_ρ_* = −0.59, *p _FDR_* = 0.004), indicating that greater preference for social touch was associated with lower FC strength during nonsocial touch observation. A similar trend was observed in social touch (*_ρ_* = −0.41, *p _FDR_*= 0.06). This relationship was absent in non-autistic individuals (Figure 7B). FC strength between these two networks did not differ across groups and touch types.

Lastly, two groups showed differences in how STQ scores affected FC between the limbic system and social perceptual network. ANOVA results showed a significant interaction between group, touch type, and STQ scores affecting FC strength (*_χ_2*(1, 42) = 11.66, *p _FDR_* = 0.009). However, FC strength did not differ across groups and touch types. Correlation analysis showed only a marginal trend of positive correlation between STQ scores and FC strength in the AUT group during nonsocial touch observation (*_ρ_* = 0.42, *p _FDR_* = 0.06). This relationship was absent during social touch in the AUT group and during both conditions in the NON-AUT group.

## Discussion

The current study investigated how brain connectivity architecture is functionally modulated during the observation of social (human-to-human) versus non-social (human-to-object) touch in AUT compared to NON-AUT. It also explored how these neural functional properties are associated with individual differences in social responsiveness and an attitude towards social touch. By applying both hypothesis- and data-driven approaches, the study found that, at the level of specific brain regions, AUT showed the increased FC between areas involved in visual, social, and somatosensory processing (Figure 4), with some of these connections being modulated by the extent to which AUT prefer social touch in their daily lives and their level of social responsiveness (Figure 5). At the large-scale brain network level, atypical connectivity was primarily observed in the sensorimotor network, where the AUT group showed either reversed context-based modulation (Figure 6) or lacked the association between FC and an attitude towards social touch, unlike the NON-AUT group (Figure 7).

### Increased connectivity between brain areas in autism and its link with the social responsiveness and an attitude towards social touch

Twelve brain areas functionally relevant to touch observation were extracted, and connectivity was measured for each ROI pair across two groups and touch conditions. The early visual and early somatosensory areas exhibited increased FC strength in AUT compared to NON-AUT under both social and nonsocial touch conditions, with observing social touch resulting in a stronger FC in both groups (Figure 4A). Consistent with previous fMRI and EEG findings (Lee Masson et al., 2020b; Lee Masson and Isik, 2023), the increased FC in response to observed social touch in both groups highlights the importance of neural communication between these early sensory areas in processing the social aspects of human touch behavior. The current finding further elucidates the role of the visuo-tactile mirroring mechanism during social touch observation (Blakemore et al., 2005; Ebisch et al., 2008; Serino et al., 2008; Bolognini et al., 2011), with the novel insight that this mechanism may be particularly heightened in autism. Additionally, within the AUT group, the strength of FC between the visual motion area and early somatosensory area was associated with individual differences in the attitude towards social touch (Figure 5B). Specifically, individuals with a more positive attitude toward social touch demonstrated increased FC during social touch observation, a relationship that was not observed in NON-AUT.

While the underconnectivity theory has been predominant in autism research, overconnectivity has also been observed across various brain regions and networks (Delmonte et al., 2013; Keown et al., 2013; Cerliani et al., 2015; Chien et al., 2015; Abbott et al., 2016; Fishman et al., 2018; Chen et al., 2021). Of these studies, those relevant to the current work examined resting FC in autistic toddlers (Chen et al., 2021). Increased resting FC between the visual and sensorimotor networks during sleep was observed in these toddlers, and this overconnectivity was linked to more severe autism. The authors suggested that such overconnectivity in sensory brain circuits may negatively impact early development, leading to the atypical behaviors observed in autism. However, as the study was not longitudinal, the specific effects of these altered connections on social information processing later in life remain unknown. In contrast to the view that overconnectivity has a negative impact on behavioral outcome, the observation that both groups showed increased FC in early sensory networks during social touch observation, along with similar results found in previous research with the NON-AUT group (Lee Masson et al., 2020b), suggests that the heightened FC between visual and somatosensory areas in AUT may serve as a compensatory neural mechanism to support social processing. The current finding also suggests that this neural strategy seems more prevalent in individuals who have a more positive attitude towards social touch within the AUT group.

Similarly, the visual and social brain regions displayed increased FC in response to social touch, compared to nonsocial touch in AUT – a difference not observed in NON-AUT (Figure 4B). This effect in AUT was linked to individual differences in social responsiveness; those with greater social responsiveness showed increased FC during both types of touch, with a more pronounced increase for social touch (Figure 5A). These findings further suggest that the neural compensatory mechanism may be more engaged when AUT process social touch events. This mechanism seems more prevalent in those who show higher levels of social responsiveness.

Emerging research has observed neural compensatory mechanisms in autism that are often accompanied by typical behavioral performance. These mechanisms involve enhanced brain responses (Lai et al., 2019; Moessnang et al., 2020), the activation of additional brain areas (Subbaraju et al., 2018), and increased connectivity (Jasmin et al., 2023). Specifically, increased FC between the inferior frontal cortex and social communication regions (i.e., middle and anterior STS) was observed during spontaneous conversations (Jasmin et al., 2023). This increase in FC was associated with more typical language behavior in AUT. Similarly, a recent study using the same touch video stimuli suggested that greater cognitive effort, as reflected in larger pupil dilation in response to social touch videos, may act as a compensatory mechanism (Lee et al., 2024). These findings highlight that, despite the common negative interpretation of overconnectivity (Maximo et al., 2014), it may, in some cases, support rather than impede social behavior in autism. However, compensatory mechanisms are not always beneficial, and some may be linked to camouflaging, which can negatively affect mental health (Field et al., 2024). Further research is needed to explore the complex relationship between neural compensation, cognitive effort, camouflaging, and mental health outcomes, particularly in relation to social touch.

### Atypical connectivity of the sensorimotor network with other brain networks implicated in social-emotional processing

To gain a comprehensive understanding of how large-scale brain networks coordinate functional communication in response to both social and nonsocial touch events across groups, and to complement the findings from the ROI-based analysis, 10 large-scale brain networks formed during touch observation were extracted using a data-driven ICA method. These networks included one visual, one executive control, one sensorimotor, one social perceptual, two salience, one limbic, one reward, and two default mode networks. Examination of FC between large-scale brain networks primarily revealed atypicality in the sensorimotor network.

In the AUT group, the sensorimotor network showed stronger connectivity with the posterior salience network during nonsocial touch. Conversely, the NON-AUT group showed stronger connectivity during social touch (Figure 6). These findings suggest that for nonsocial touch – typically perceived as having less emotional significance (i.e., more neutral and less arousing (Lee Masson and Op de Beeck, 2018)) – the AUT group exhibits heightened coordination between the systems implicated in embodied simulation and affective interoception. This finding extends prior research by revealing the (absent) involvement of embodied simulation in social versus nonsocial touch observation and aligns with the systemic literature showing atypical embodied simulation for emotional stimuli, while remaining intact for non-emotional stimuli (Hamilton, 2013).

Prior research has identified similar patterns in AUT with respect to receiving touch (Kaiser et al., 2016). Autistic children and adolescents show reduced activity in brain areas implicated in social-emotional processing, including the insula, MTG, and STG – parts of the posterior salience network identified in this study (Figure 3 and Table S1) – when receiving affective touch. In contrast, they show increased activity in the primary somatosensory cortex and the insula when receiving non-affective touch. The current findings may be partially attributed to sensory reactivity differences associated with autism. Some nonsocial video clips used in this study feature a person exploring clothing with various textures, which may have triggered hyper-reactivity and sensory-seeking in autism (MacLennan et al., 2022). Further research is needed to explore the relationship between sensory-seeking and heightened network communication during the observation of human-object interactions, with objects that individuals find particularly engaging.

The sensorimotor network identified in this study primarily comprises the precentral and postcentral gyri, while the posterior salience network includes the insula, middle and anterior regions of the MTG, STS, and STG (Figure 3, Table S1). Although temporal regions are less commonly associated with the posterior salience network, they may work in conjunction with the insula because of their role in processing a wide variety of social stimuli (Deen et al., 2015; Yang et al., 2015; Santavirta et al., 2023; Lee Masson et al., 2024). The insula, in turn, is essential for detecting the emotional significance of stimuli and for interoceptive awareness triggered by those stimuli (Augustine, 1996; Carr et al., 2003; Cauda et al., 2011; Zaki et al., 2012). This complementary relationship between the temporal regions and the insula may result in their integration into a unified network essential for processing the emotional significance of observed touch and related bodily sensations. Furthermore, both regions are often involved in processing affective touch, whether it is experienced or observed, with their neural activity influenced by the perceived pleasantness (Morrison et al., 2011; Gordon et al., 2013; Davidovic et al., 2017; Kirsch et al., 2020). The current finding is consistent with earlier research that identifies atypical insula activity and connectivity in autism (Ebisch et al., 2011b; Francis et al., 2019; Nomi et al., 2019). The current findings further contribute to our understanding by demonstrating how the connectivity of the posterior salience network – primarily involving the insula – is differently modulated based on touch type in autism.

Extensive neuroimaging research has demonstrated somatosensory activation in response to observed social touch, highlighting the vital role of embodied simulation in understanding another person’s touch experiences (Blakemore et al., 2005; Pihko et al., 2010; Gazzola et al., 2012; Gallese and Ebisch, 2013; Lee Masson et al., 2018; Schirmer and McGlone, 2018; Schaefer et al., 2024). The degree of this activation has been associated with autism and other individual differences, including social touch preference and the empathy (Schaefer et al., 2012; Bolognini et al., 2014; Peled-Avron et al., 2016; Lee Masson et al., 2018, 2019). Reduced embodied simulation has also been observed in other social cognitive domains in relation to autism. Research has shown that AUT use a visuo-spatial strategy for perspective-taking tasks, while NON-AUT rely more on embodied simulation (Conson et al., 2015). Reduced somatosensory responses have been also found in autism during facial expression discrimination tasks (Fanghella et al., 2022). The vital role of embodied simulation in understanding another person’s touch experiences is further emphasized by the current finding, which shows that during social touch observation, NON-AUT with a positive attitude towards social touch exhibit greater functional coordination between the sensorimotor system and the DMN – a network implicated in social cognition (Mars et al., 2012; Li et al., 2014) (Figure 7A). In contrast, in autism, this functional coordination between the two networks associated with embodied simulation and social cognition does not correlate with social touch preference.

Other factors might explain the differences in network connectivity. AUT experience social touch differently, including which body parts they find pleasant or appropriate (Mello et al., 2024). Since the social video clips used in the current study feature social touch between NON-AUT, the atypical connectivity in the embodied simulation system may be partially explained by this difference. When people with differing experiences interact, they may struggle to empathize (Milton, 2012) and rely more on cognitive strategies rather than embodied simulation, a phenomenon also observed in the source study with multivoxel pattern analysis (Lee Masson et al., 2019). However, the current study does not include an experimental condition in which NON-AUT observe social touch interactions involving AUT. Similarly, factors other than autism, such as alexithymia and anxiety, might also partially explain the atypical connectivity. These co-occurring conditions associated with autism are reported to impact embodied simulation, emotion recognition, and the ability to empathize with others (Lassalle et al., 2019; Raman et al., 2023). Further research is needed to address these questions.

### Converging evidence from both ROI-based and ICA-based analyses of connectivity in autism

These two approaches complement each other by providing insights into neural mechanisms that one alone cannot reveal. The ROI approach sheds light on connectivity between specific brain regions, while the ICA-based FC approach, using data-driven methods, offers additional evidence at the network level that the ROI method might overlook. Some discrepancies emerged between the results obtained from the two methodologies. Findings from brain region analyses reveal hyper-connectivity between the early visual areas, early somatosensory regions, and social brain areas, potentially linked to neural compensatory mechanisms. In contrast, large-scale brain network analyses reveal atypical connectivity between networks involved in embodied simulation, affective interoception, and social cognition, potentially linked to sensory reactivity and qualitative variations in social touch experiences between AUT and NON-AUT. Further research is needed to explore why neural compensatory mechanisms seem to occur only in brain areas, while the network predominantly exhibits connectivity differences. Importantly, none of the current findings can be attributed to head motion, as both groups exhibited similar magnitudes of head motion. Moreover, outlying volumes with higher framewise displacement were removed, and motion parameters were included as covariates in the analysis.

### Limitations

This study focused on a specific subgroup of autistic male adults with average to above-average intelligence, no language or learning difficulties, and minimal support needs. While the homogeneity of this sample helps control for variables such as age, IQ, and gender, future research would benefit from including a more diverse autistic sample. This may include females and individuals with greater support needs, such as those with below-average IQ, who may struggle to utilize compensatory cognitive strategies, potentially leading to different connectivity patterns. In particular, including a female sample is crucial, as autistic females experience touch differently compared to both autistic males and non-autistic females (Mello et al., 2024). Lastly, the current study includes only small number of participants (N=42). However, the effect sizes observed in this study are generally considered medium to large (e.g., *_ρ_* = −0.68), though the wide confidence intervals suggest some variability in these estimates.

### Conclusions

The current study presents distinct FC patterns in autism during the observation of social touch interactions and nonsocial human-object manipulation. Heightened connectivity in the sensory and social brain areas suggests a compensatory neural mechanism that supports social processing, particularly in AUT with greater social responsiveness and a more positive attitude toward social touch. These findings question the common negative view of hyper-connectivity in autism, proposing that such connectivity might sometimes serve as an adaptive strategy. At the network level, connectivity differences between the sensorimotor, salience, and default mode networks points to altered information flow, suggesting that the engagement of embodied simulation, affective interoception, and social cognition during touch observation may differ in autism. The increased connectivity of the sensorimotor network during nonsocial touch highlights that the extent of embodied simulation varies with the type of touch and is not always absent in autism. In terms of methodology, this study highlights the advantages of using a task-based connectivity approach to examine context-sensitive changes in neural functional architecture in autism, providing evidence that a task-free resting state approach may not offer. Finally, this study links neural measures to behaviors, offering insights into how brain connectivity in response to observed touch relates to autistic social behavior and preferences for social touch.

## Availability of data and materials

Data that support the findings of this study are available through the Open Science Framework (https://osf.io/yhvjd/) for scientific use.

## Acknowledgements

The author thanks Dr Deborah Riby and Dr Jeyoung Jung for discussion and feedback on the manuscript.

