## Supplementary Table 1 for "Dynamic functional adaptations during touch observation in autism: Connectivity strength is linked to attitudes towards social touch and social responsiveness"

**Supplementary Materials.**

Table S1. **The implicated brain regions and MNI coordinates for the networks. Cluster size > 200 voxels; P _FWE_ < 0.001.**

| **Network** | **Component description** |  | **Area** | **Peak X** | **Peak Y** | **Peak Z** | ***T* Statistics** | **N Voxels** |
| --- | --- | --- | --- | --- | --- | --- | --- | --- |
| IC1 | Higher Visual Network | R | superior occipital gyrus | 30 | -88 | 11 | 15.45 | 2498 |
| IC4 | Sensorimotor Network | L | precentral gyrus medial segment | -3 | -19 | 62 | 17.80 | 2760 |
| IC11 | Posterior Salience Network | L | anterior insula | -42 | 8 | -13 | 16.02 | 628 |
|  |  | R | temporal pole | 48 | 11 | -13 | 13.46 | 479 |
| IC18 | Limbic System | R | amygdala | 27 | -4 | -19 | 37.39 | 466 |
|  |  | L | middle temporal gyrus | -54 | -7 | -19 | 15.38 | 202 |
| IC19 | Default Mode Network 1 | L | precuneus | -3 | -61 | 20 | 23.1 | 1595 |
|  |  | R | fusiform gyrus | 24 | -40 | -13 | 15.16 | 213 |
|  |  | L | superior frontal gyrus | -24 | 5 | 59 | 15 | 380 |
|  |  | R | middle occipital gyrus | 42 | -70 | 35 | 14.7 | 250 |
|  |  | L | middle occipital gyrus | -39 | -76 | 32 | 13.19 | 236 |
|  |  | R | superior frontal gyrus | 24 | 11 | 53 | 12.82 | 377 |
| IC24 | Anterior Salience Network | L | supplementary motor cortex | -6 | 14 | 50 | 18.97 | 1839 |
|  |  | L | anterior insula | -42 | 17 | -13 | 12.01 | 228 |
|  |  | L | middle frontal gyrus | -27 | 41 | 38 | 11.43 | 313 |
| IC25 | Left Executive Control Network | L | superior frontal gyrus | -15 | 35 | 47 | 16.01 | 2425 |
|  |  | L | angular gyrus | -45 | -67 | 32 | 15.39 | 1022 |
|  |  | L | middle temporal gyrus | -57 | -46 | -7 | 12.00 | 230 |
| IC26 | Reward System | L | caudate | -15 | 20 | -4 | 20.38 | 3385 |
| IC27 | Default Mode Network 2 | L | anterior cingulate gyrus | -3 | 44 | 11 | 20.09 | 2747 |
|  |  | L | posterior cingulate gyrus | -3 | -52 | 26 | 17.72 | 661 |
|  |  | R | frontal operculum | 48 | 20 | -7 | 13.10 | 353 |
|  |  | L | angular gyrus | -51 | -64 | 35 | 12.96 | 223 |
|  |  | L | inferior frontal gyrus | -54 | 20 | 8 | 12.57 | 278 |
|  |  | R | postcentral gyrus | 45 | -25 | 44 | 10.03 | 231 |
| IC28 | Social Perceptual Network | R | superior temporal gyrus | 66 | -46 | 17 | 13.09 | 892 |
|  |  | L | superior temporal gyrus | -57 | -31 | 5 | 11.84 | 712 |
